# Motor learning and transfer: from feedback to feedforward control

**DOI:** 10.1101/845933

**Authors:** Rodrigo S. Maeda, Paul L. Gribble, J. Andrew Pruszynski

**Affiliations:** Brain and Mind Institute, Western University, London, Ontario, Canada; Robarts Research Institute, Western University, London, Ontario, Canada; Dept. of Psychology, Western University, London, Ontario, Canada; Dept. of Physiology and Pharmacology, Western University, London, Ontario, Canada

**Keywords:** feedback control, internal model, intersegmental dynamics, reaching, motor learning, stretch reflex

## Abstract

Previous work has demonstrated that when learning a new motor task, the nervous system modifies feedforward (ie. voluntary) motor commands and that such learning transfers to fast feedback (ie. reflex) responses evoked by mechanical perturbations. Here we show the inverse, that learning new feedback responses transfers to feedforward motor commands. Sixty human participants (34 females) used a robotic exoskeleton and either 1) received short duration mechanical perturbations (20 ms) that created pure elbow rotation or 2) generated self-initiated pure elbow rotations. They did so with the shoulder joint free to rotate (normal arm dynamics) or locked (altered arm dynamics) by the robotic manipulandum. With the shoulder unlocked, the perturbation evoked clear shoulder muscle activity in the long-latency stretch reflex epoch (50-100ms post-perturbation), as required for countering the imposed joint torques, but little muscle activity thereafter in the so-called voluntary response. After locking the shoulder joint, which alters the required joint torques to counter pure elbow rotation, we found a reliable reduction in the long-latency stretch reflex over many trials. This reduction transferred to feedforward control as we observed 1) a reduction in shoulder muscle activity during self-initiated pure elbow rotation trials and 2) kinematic errors (ie. aftereffects) in the direction predicted when failing to compensate for normal arm dynamics, even though participants never practiced self-initiated movements with the shoulder locked. Taken together, our work shows that transfer between feedforward and feedback control is bidirectional, furthering the notion that these processes share common neural circuits that underlie motor learning and transfer.

## Introduction

Previous studies have demonstrated that learning a new motor task modifies feedforward (i.e. voluntary) motor commands and that such learning also changes the sensitivity of fast feedback responses (i.e. reflexes) to mechanical perturbations (Wang et al., 2001; Wagner and Smith, 2008; Ahmadi-Pajouh et al., 2012; Yousif and Diedrichsen, 2012; Cluff and Scott, 2013; Maeda et al., 2018, 2019). For example, after people learn to generate straight reaching movements in the presence of an external force field or learn to reduce shoulder muscle activity when generating pure elbow movements with shoulder fixation, evoked stretch reflex responses to mechanical perturbations reflect (Ahmadi-Pajouh et al., 2012; Maeda et al., 2018, 2019) and correlate with (Cluff and Scott, 2013) the learning expressed during self-initiated reaching. Such transfer from feedforward motor commands to feedback responses is thought to take place because of shared neural circuits at the level of the spinal cord, brainstem and cerebral cortex (Pruszynski, 2014; Scott, 2016).

The presence of shared neural resources also predicts transfer from feedback responses to feedforward motor commands. However, little is known about such transfer presumably because it is relatively hard to elicit learning in reflexes without engaging associated voluntary responses following mechanical perturbations. Here we tested for such transfer by leveraging two approaches to elicit stretch reflex responses while minimizing engagement of voluntary motor commands in the learning process. First, we used mechanical perturbations with very short durations (20 ms) previously shown to elicit reflex responses but very weak or non-existent voluntary muscle responses (Ghez and Shinoda, 1978; Lee and Tatton, 1982; Lewis et al., 2005; Schuurmans et al., 2009; Kurtzer et al., 2010; Kurtzer, 2019). Second, we instructed participants to not intervene with the mechanical perturbations previously shown to substantially reduce or eliminate voluntary muscle responses (Asatryan and Feldman, 1965; Crago et al., 1976; Calancie and Bawa, 1985; Shemmell et al., 2009; Forgaard et al., 2015, 2016, 2019).

Human participants received short-duration mechanical perturbations (20 ms) at their shoulder and elbow balanced in a way that yielded pure elbow motion (Kurtzer et al., 2008, 2014; Maeda et al., 2017, 2018; Kurtzer, 2019). Perturbations were applied with the shoulder joint free to rotate (normal arm dynamics) or with the shoulder locked by the robotic manipulandum (novel arm dynamics). Locking the shoulder joint alters the mapping between joint torques and joint motion because torques that normally arise at the shoulder joint due to elbow rotation are cancelled at the robotic apparatus. We report two key findings. First, we found that shoulder reflex responses were gradually reduced on a timescale of hundreds of trials as appropriate for the novel arm dynamics. Second, we found that this reduction transferred to voluntary motor commands as we observed 1) a reduction in shoulder extensor muscle activity during self-initiated elbow reaching trials, even though participants never practiced reaching movements with the shoulder locked and 2) kinematic errors (ie. aftereffects) after releasing the shoulder joint again in the direction predicted when failing to compensate for normal arm dynamics. Taken together, our work furthers the idea feedforward and feedback control share common neural circuits (Wagner and Smith, 2008; Ahmadi-Pajouh et al., 2012; Cluff and Scott, 2013; Maeda et al., 2018) by demonstrating that transfer between voluntary motor commands and reflex responses is bidirectional.

## Materials and Methods

### Subjects

Sixty healthy human participants (aged 17–39, 34 females) took part in this study. Participants self-reported that they were right-handed and free from visual, neurological, or musculoskeletal deficits. All participants were naive as to the purpose of the study, were free to withdraw at any time, and provided written informed consent before participating. The Office of Research Ethics at Western University approved this study.

### Apparatus

Participants performed the experiments with a robotic exoskeleton (KINARM, Kingston, ON, Canada). As described in previous studies (Scott, 1999; Pruszynski et al., 2008, 2009), this device allows for flexion and extension movement of the shoulder and elbow joints in the horizontal plane, and can independently apply torque loads at these joints. Target lights and hand cursor feedback cursor were presented in the same plane as the movement using an overhead LCD monitor and a semi-silvered mirror. Direct vision of the arm was occluded with a physical shield. To ensure a comfortable and tight coupling between each participant’s arm and the robot, the two segments of the exoskeleton robot (upper arm and forearm) were adjusted for each participant arm and the spaces were filled with a firm foam. Lastly, the robot was calibrated so that the projected hand cursor was aligned with each participant’s right index finger.

### Experimental task and protocols

Twenty participants used a robotic apparatus and received 20 ms torque pulse perturbations that caused pure elbow motion with the shoulder joint free to move and with the shoulder fixed by the robotic manipulandum (altered arm dynamics). Before and after learning with the shoulder fixed participants performed twenty-degree elbow extension movements (probes).

In the beginning of a trial, participants were instructed to keep their hand in a home target (white circle, 0.6 cm diameter) which required shoulder and elbow angles of 40° and 80° (external angles), respectively (Figure 1A). After a random period (250-500 ms, uniform distribution), a background load (+2 Nm) was slowly introduced (rise time = 500ms) to the elbow joint to ensure baseline activation of shoulder and elbow extensor muscles (Maeda et al., 2018). After an additional random hold period (1-2 s, uniform distribution), a step-torque of 20 ms duration (i.e., perturbation) was applied to the shoulder and elbow joints (+/-2 Nm at each joint over and above the background torque). At the same time, the home target and hand feedback were turned off and participants were instructed to not intervene the perturbation. Critically, we chose this combination of shoulder and elbow loads to minimize shoulder motion (see Kurtzer et al., 2008, 2014; Maeda et al., 2017; Kurtzer, 2019). Within a random period (300-500 ms, uniform distribution), a servo-controller brought the participant’s arm back to the home location, and the same procedure was repeated for a new trial. The servo-controller was implemented as a stiff, viscous spring and damper (K = 500 N/m and B = 250 N/(m/s)) with ramp up time of 100ms and trajectory duration of 500ms. Again, participants were instructed to not intervene with the servo (Figure 1B, Top left column).

**Figure 1:**
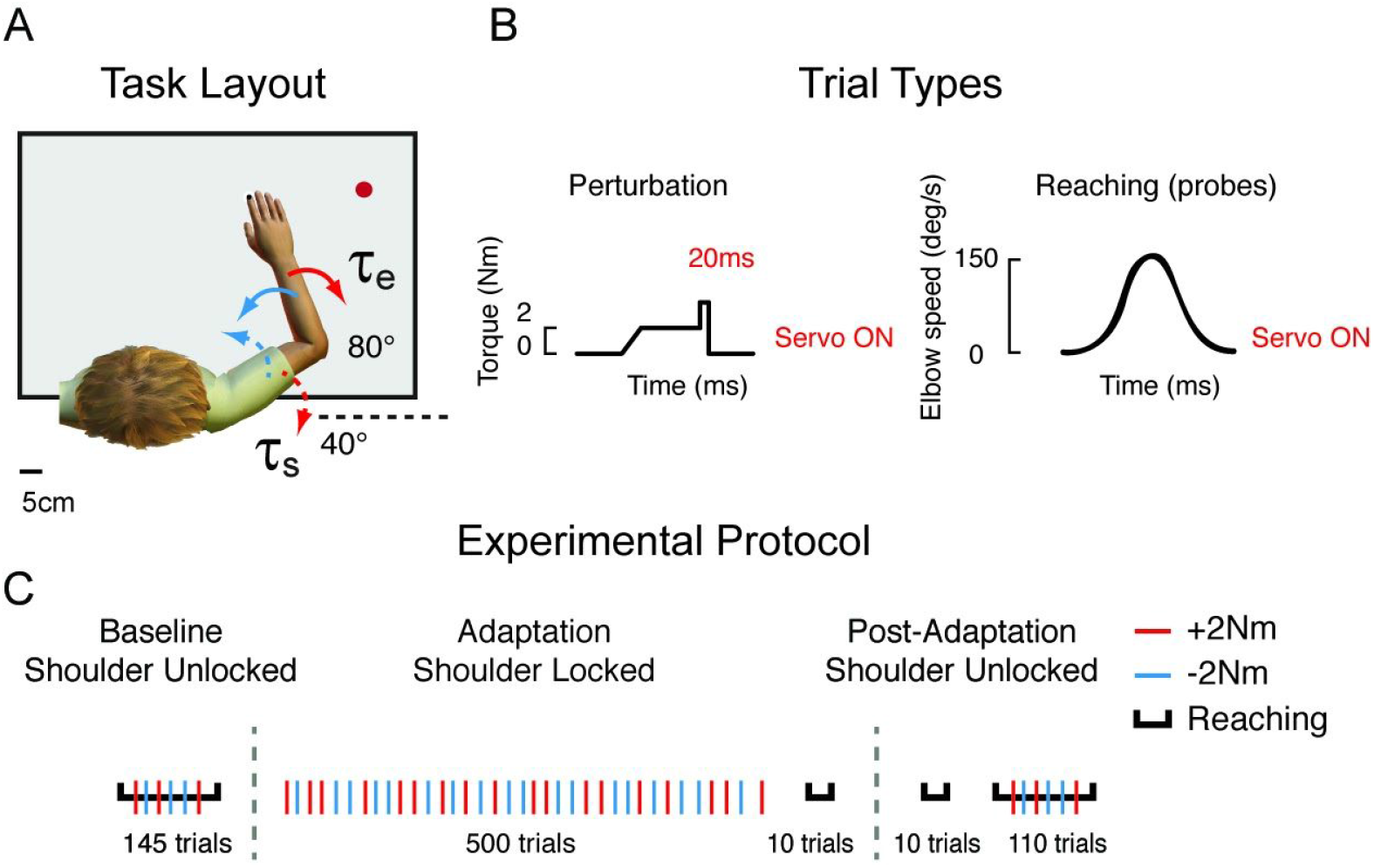
Experimental setup. **A**, Participants either received mechanical perturbations that created pure elbow motion (learning) or generated pure elbow extension movements to a dark red goal target (probes). Red and blue arrows represent the direction of the multi-joint step-torques applied to the shoulder and elbow joints. **B**, These perturbations lasted 20 ms (load duration) and reaching trials had speed constraints (100-180ms duration). **C**, Illustration of the experimental protocol. Participants performed 145 baseline trials with the shoulder joint unlocked (learning and probes), 500 adaptation trials with the shoulder joint locked (learning trials), 10 probe trials with the shoulder locked, 10 probe trials with the shoulder unlocked and 110 post-adaptation trials with the shoulder joint unlocked (learning and probes). Black horizontal lines lines indicate blocks with reaching trials and red and blue lines indicate trials with the direction of the multi-joint step-torques as shown in **A**.

In some trials, participants were required to perform reaching movements (probes). Reaching trials occurred before learning with the shoulder unlocked, after learning with the shoulder locked, and after learning with the shoulder unlocked. In these trials, participants started by also keeping their hand in the same home target (red circle, 0.6 cm diameter). After a random hold period (250 –500 ms, uniform distribution), a goal target (white circle: 3 cm diameter) was presented in a location that could be reached with a 20° pure elbow extension movement. Participants were required to remain at the home location for an additional random period (250 – 500 ms, uniform distribution) so that the goal target turned red, the hand feedback cursor was turned off and participants were allowed to start the movement. Participants were instructed to move to the goal target with a specific movement speed. The goal target turned green when movement time (from exiting the home target to entering the goal target) was between 100 and 180 ms, orange when it was too fast (<100 ms) and red when it was too slow (>180 ms). No restrictions were placed on movement trajectories. Participants were required to remain at the goal target for an additional 500 ms to finish a trial. The hand feedback cursor turned back on when the participant’s hand entered the goal target or after a fixed time window of 500ms from the cue to start of the movement (Figure 1B, top right column). After a random period (300-500 s, uniform distribution), the servo-controller, as described above, moved the participant’s arm back to the home location. In five percent of reaching trials, the background torques turned on, remained on for the same time period (1.0–2.5 s, uniform distribution), but then slowly turned off, after which participants were still required to perform the reaching movements. These trials ensured that background loads were not predictive of perturbation trials. The order of all perturbation and reaching trials was randomized in the baseline and post adaptation phases and all perturbations were randomized in the adaptation phase.

Participants first completed a total of 145 baseline trials (120 mechanical perturbations and 25 reaching, randomized), with the shoulder joint free to move. We then locked the shoulder joint with a physical clamp, and participants completed 500 perturbation trials (adaptation phase) and 10 reaching trials (probes) with the shoulder joint locked. We then unlocked the shoulder joint and participants completed 10 reaching movements with the shoulder joint unlocked. Lastly, participants completed a total of 110 post-adaptation trials (100 mechanical perturbations and 10 reaching, randomized), (post-adaptation phase) (Figure 1C).

Ten additional participants performed the same version of this experiment but without locking the shoulder joint. This served as a control to rule out changes that could be caused by extensive exposure to perturbations rather than learning associated with the shoulder fixation manipulation.

Twenty additional participants performed the same experimental procedures as above with the exception of the duration of the mechanical perturbation which lasted 100 ms. This served to quantify and contrast the amount of learning and transfer that would be observed when the perturbation evoked substantial activity in the voluntary response epoch. Lastly, ten participants (control) performed the same version of this Experiment (with mechanical perturbations of 100ms load duration) but without locking the shoulder joint, which served as its associated control experiment for exposure.

All experiments lasted about 2h. Rest breaks were given throughout or when requested. Before starting, participants completed practice trials until they expressed that they understood the instructions and comfortably achieved ~90% success in reaching trials (approx. 10 min).

### Kinematic recordings and analysis

We recorded movement kinematics (i.e. hand position, and joint angles) with the robotic device at 1000 Hz and then low-pass filtered offline (12 Hz, 2-pass, 4th-order Butterworth). In Experiments 1 and 2, data from perturbation trials was aligned on perturbation onset and data from reaching trials was aligned on movement onset, defined as 5% of peak elbow angular velocity (see Gribble and Ostry, 1999; Maeda et al., 2017, 2018). We quantified aftereffects of reaching movements following shoulder fixation by calculating hand path errors relative to the center of the target at 80% of the movement between movement onset and offset (also defined at 5% from the peak angular elbow velocity). This window was used to select the kinematic traces before any corrections (Maeda et al., 2018).

### EMG recordings and analysis

Electromyographic signals (EMG) were amplified (gain = 10^3^) and digitally sampled at 1000 Hz (Delsys Bagnoli-8 system with DE-2.1 sensors, Boston, MA). EMG surface electrodes were used and placed on the skin surface on top of the belly of five upper limb muscles (pectoralis major clavicular head, PEC, shoulder flexor; posterior deltoid, PD, shoulder extensor; biceps brachii long head, BB, shoulder and elbow flexor, Brachioradialis, BR, elbow flexor; triceps brachii lateral head, TR, elbow extensor). Before electrode placement, the participant’s skin was prepared with rubbing alcohol, and the electrodes were coated with conductive gel. Electrodes were placed along the orientation of muscle fibers. A reference electrode was placed on the participant’s left clavicle. EMG data were band-pass filtered (20–500 Hz, 2-pass, 2nd-order Butterworth) and full-wave rectified offline.

First, we investigated whether feedback responses in shoulder muscles adapt to a novel arms dynamics following shoulder fixation. To assess whether the short and long latency stretch response of shoulder and elbow extensor muscles account for and adapt over time to novel arm’s dynamics, we binned the EMG data into previously defined epochs (see Pruszynski et al., 2008). This included a pre-perturbation epoch (PRE, –50-0 ms relative to perturbation onset), the short-latency stretch response (R1, 20-50 ms), the long-latency stretch response (R2/3, 50-100 ms), and the voluntary response (VOL, 100-150ms).

We also investigated whether shoulder and elbow muscles during pure elbow reaching adapt after learning novel arm dynamics following shoulder fixation during perturbation trials. To compare the changes in amplitude of muscle activity before and after learning, we calculated the mean amplitude of phasic muscle activity in a fixed time-window, −100 ms to +100 ms relative to movement onset (see Debicki and Gribble, 2005; Maeda et al., 2017, 2018). These windows were chosen to capture the agonist burst of EMG activity in each of the experiments.

Data from perturbation trials were normalized by the pre-perturbation activity, which was the activity required to compensate for a 2 Nm constant load. Data from reaching trials were normalized by a pre-activity, which was the activity required to compensate for a 1 Nm constant load in normalization trials performed prior to the experiments (see Pruszynski et al., 2008; Maeda et al., 2017, 2018). Data processing was performed using MATLAB (r2018a, Mathworks, Natick, MA).

### Statistical analysis

Statistical analyses were performed using R v3.2.1 (Boston, MA). We performed different statistical tests (e.g., repeated measures ANOVA with Tukey tests for multiple comparisons, t-test, and regression analysis), when appropriate in each of the two experiments. Further details of these analyses are provided in the Results. Experimental results were considered statistically significant if the corrected p-value was less than < 0.05.

## Results

### Reducing feedback responses with shoulder fixation

Participants (N=20) held their hand in a home target and mechanical perturbations were applied to their arm. The mechanical perturbations were 20 ms duration torque pulses applied simultaneously to the shoulder and elbow joints which yielded minimal shoulder motion but different amounts of elbow motion (Figure 2A). In selected trials (probes) participants voluntarily moved their hand between two targets that were placed at locations that required 20° of pure elbow extension.

**Figure 2:**
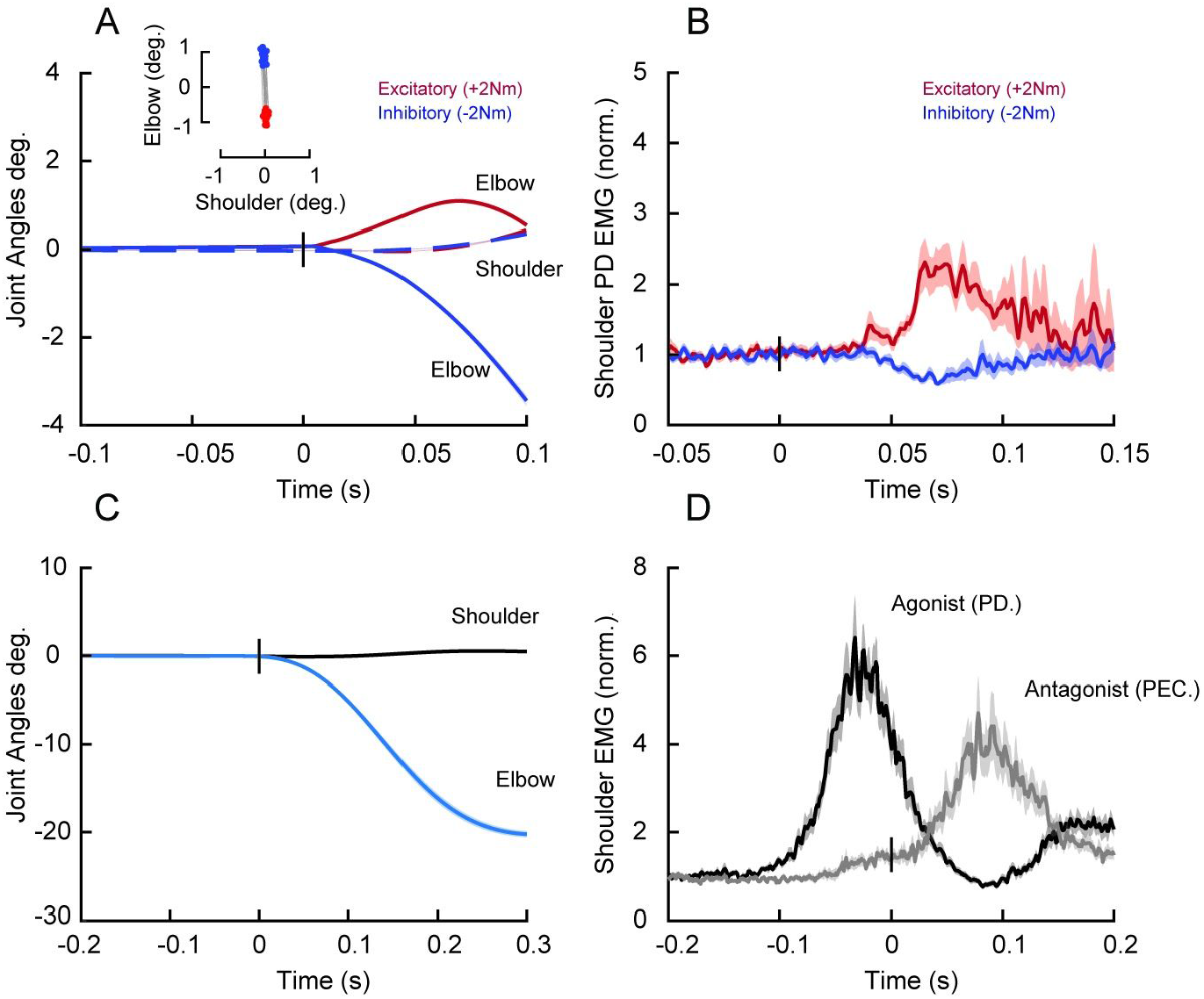
Compensating for intersegmental dynamics during perturbations that created pure elbow motion and when generating pure elbow extension movements. **A**, Average kinematics of the shoulder (dashed) and elbow (solid) joints following mechanical perturbations of 20 ms duration. Red and blue traces are from the shoulder/elbow flexor torque and shoulder/elbow extensor torque conditions, respectively. Shaded areas represent the standard error of the mean. Data are aligned on perturbation onset. Inset shows the amount of shoulder and elbow displacement 50 ms post-perturbation (data are shown for all subjects). **B**, Normalized shoulder muscle activity associated with panel **A. C**, Average kinematics of the shoulder (black traces) and elbow (blue traces) joints for elbow extension trials. Shaded areas represent the standard error of the mean. Data are aligned on movement onset. D, Black and gray lines represent average agonist (PD) and antagonist (PEC) muscle activity associated with the movement in **C**.

As previously demonstrated, the mechanical perturbations elicited shoulder long-latency stretch reflexes, a response that is appropriate for countering the underlying joint torques (Figure 2A-B) (Soechting and Lacquaniti, 1988; see Kurtzer et al., 2008, 2009, 2014; Pruszynski et al., 2011; Maeda et al., 2017, 2018; Kurtzer, 2019). Also as previously demonstrated, even though they were not explicitly instructed to do so, participants performed the voluntary reaching trials by almost exclusively rotating the elbow joint, which generated substantial shoulder muscle activity as required to compensate for interaction torques that arise at the shoulder when the forearm rotates about the elbow joint (Figure 2C-D) (Gribble and Ostry, 1999; Maeda et al., 2017, 2018).

We first tested whether the nervous system adapts stretch reflex responses to the altered intersegmental dynamics following shoulder fixation by reducing shoulder muscle responses to mechanical perturbations. Figure 3A illustrates mean shoulder extensor muscle activity in the long-latency epoch (50-100 ms post-perturbation, see Methods) across trials before (i.e. baseline trials), during (i.e. adaptation trials) and after (i.e. post-adaptation trials) the shoulder joint was physically locked. Muscle activity in the long-latency epoch slowly decreased over the course of the adaptation trials and quickly returned to baseline after the shoulder lock was removed (Figure 3A). A one-way ANOVA comparing shoulder extensor muscle activity in the long-latency epoch late (last 10 trials) in the baseline, adaptation and post-adaptation phases revealed a reliable effect of phase on shoulder muscle responses (F_2,38_ = 10.92, P < 0.0001, Figure 3B-C). Tukey post-hoc tests showed that shoulder extensor muscle activity in the long-latency epoch decreased by 47% relative to baseline (P < 0.0001) after learning with the shoulder locked, and then returned to levels indistinguishable from baseline after the shoulder was unlocked (P = 0.87; Figure 3C). We found no corresponding changes in the elbow TR muscle responses in the long-latency epoch (one-way-ANOVA, = 1.244, P = 0.3, Figure 3D-F). We also found no corresponding changes in shoulder and elbow extensor muscles in the voluntary response (100-150 ms post-perturbation, see Methods) to perturbations following shoulder fixation (one-way-ANOVA in shoulder PD muscles, F_2,38_ = 1.14, P = 0.32, Figure 4 A-B and elbow TR muscles, F_2,38_ = 4.091, P = 0.02, Tukey post-hoc confirmed no reliable changes between baseline and adaptation phases, p = 0.052, see Figure 4 C-D).

**Figure 3:**
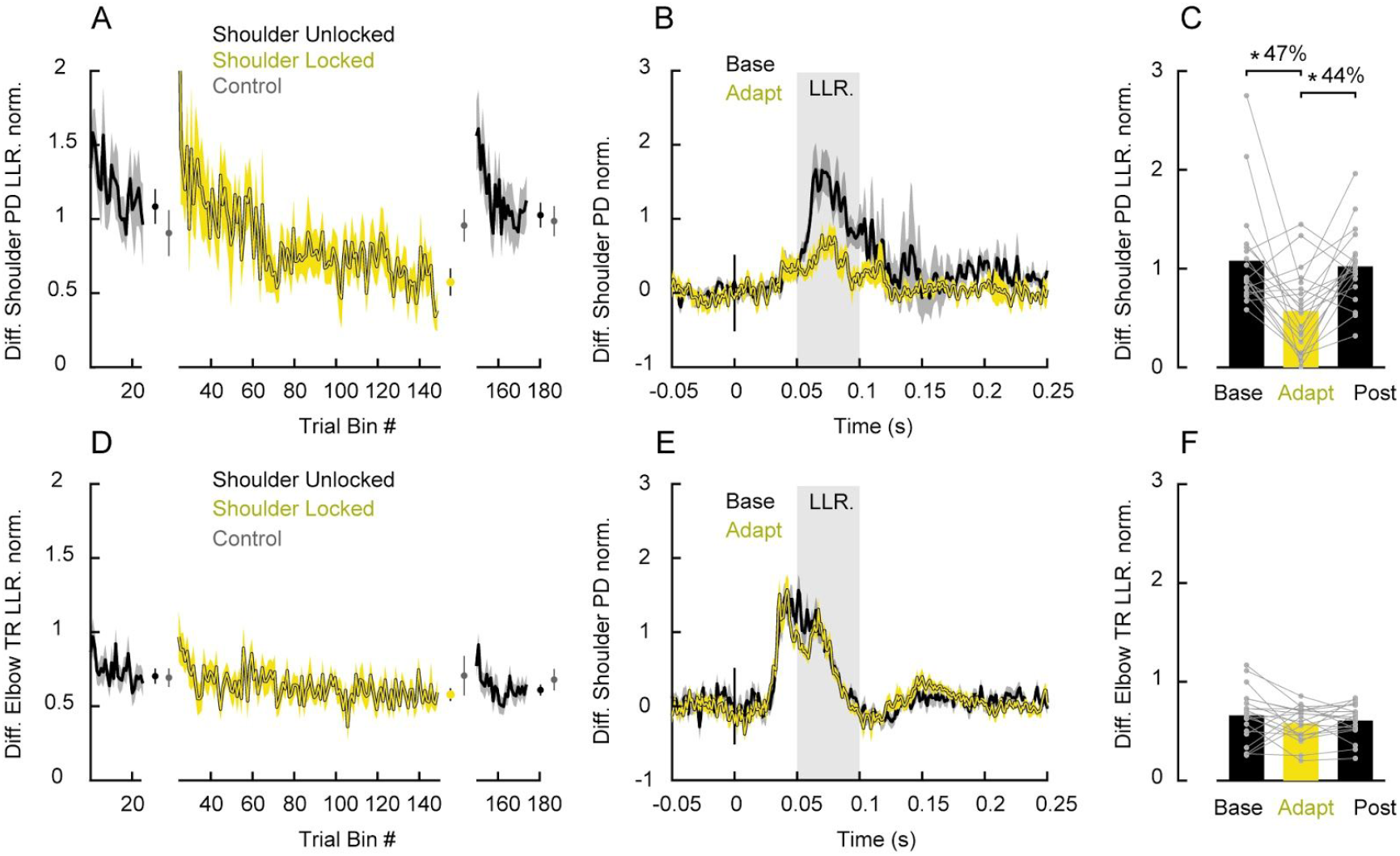
Learning novel arm dynamics by long-latency stretch reflexes during perturbation trials with shoulder fixation. **A**, Average of the difference of shoulder PD muscle activity (excitatory - inhibitory) in the long latency epoch (50-100ms post perturbation) across trials. Each data bin is 2 trials. Shaded areas represent the standard error of the mean. EMG data are normalized as described in the Methods. Error bars plotted between the phases represent the mean and standard error of the last 5 bins of trials in each phase contrasted with the respective bins in the control experiment in gray. **B**, Time series of the difference of PD normalized muscle activity averaged over the last 10 baseline and adaptation trials. Shaded areas represent the standard error of the mean. Data is aligned on perturbation onset. **C**, Average PD muscle activity in the long-latency epoch associated with trials late in the baseline, adaptation, and post-adaptation. Each dot represents data from a single participant. Asterisks indicate reliable effects (p < 0.05, see main text). **D-F**, data for the difference of elbow TR muscle (excitatory-inhibitory) are shown using the same format as **A-C**.

Importantly, we investigated whether there was a change in baseline EMG activity prior to perturbation onset across phases, which could potentially explain these changes in EMG in the long-latency epoch via the known sensitivity of this response to the pre-perturbation state of the motor neuron pool (gain scaling, Marsden et al., 1976; Bedingham and Tatton, 1984; Pruszynski et al., 2009). A one-way ANOVA comparing the activity of shoulder extensor muscle activity in the pre-perturbation epoch (−50-0 ms) as a function of experimental phase revealed no reliable effect (F_2,38_ = 1.67, P = 0.20) suggests that this mechanisms is not the driver of reflex sensitivity.

We also performed a control experiment to rule out the possibility that the reflex changes we observed above merely reflect extensive exposure to perturbations rather than learning associated with the shoulder fixation manipulation per se. Participants (N = 10) performed the same number of trials but never experienced the shoulder locked condition. Our data indicated this is not the case. We found no reliable decrease in shoulder muscle responses in the long-latency epoch across experimental phase (F_2,18_ = 0.27, P = 0.76; see gray error bars in Figure 3A). Moreover, shoulder muscle activity at the end of the adaptation phase in the main experiment (i.e. with the shoulder locked) was smaller than at the equivalent point in this control experiment (t_11_ −2.51, P = 0.023).

**Figure 4:**
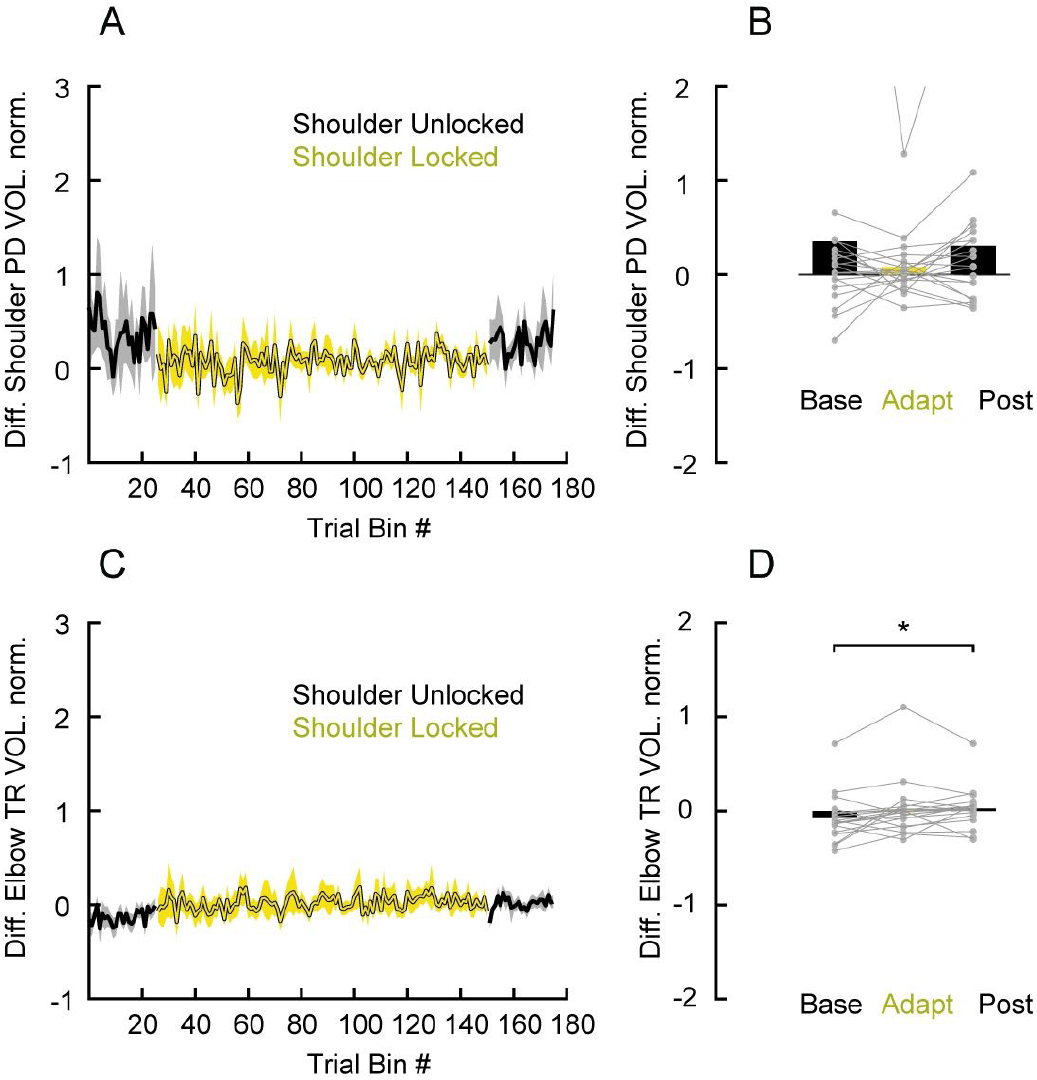
No effects of learning in the voluntary response to perturbation trials of 20 ms duration with shoulder fixation. **A**, Average of the difference of shoulder PD muscle activity (excitatory - inhibitory) in the voluntary epoch (100-150ms post perturbation) across trials. Each data bin is 2 trials. Shaded areas represent the standard error of the mean. EMG data are normalized as described in the Methods. **B**, Average PD muscle activity in the voluntary epoch associated with trials late in the baseline, adaptation, and post-adaptation. Each dot represents data from a single participant. Asterisks indicate reliable effects (p < 0.05, see main text). **C-D**, data for the difference of elbow TR muscle activity (excitatory-inhibitory) are shown using the same format as **A-B**.

### Transfer to reaching

We then investigated whether the learning exhibited by long-latency stretch reflexes during postural perturbations transferred to self-initiated reaching. Figure 5A illustrates the average time series data of shoulder extensor muscle activity in probe trials before learning (i.e. baseline trials) and after experiencing shoulder fixation during perturbations trials (i.e. adaptation trials). Figure 5B shows the mean shoulder extensor muscle activity, in a fixed time window (−100 to 100 ms, see Methods) relative to movement onset across trials before learning. Consistent with transfer, a one-way ANOVA comparing shoulder extensor muscle activity in the baseline, adaptation and post-adaptation phases (10 trials) revealed an effect of phase on shoulder muscle activity (F_2,38_ = 13.01, P < 0.0001). Tukey post-hoc tests revealed that shoulder extensor muscle activity decreased by 24% relative to baseline (P < 0.0001) and then reliably returned to baseline levels after unlocking the shoulder joint (P = 0.65; Figure 5A-B). We found no corresponding changes in elbow extensor muscle activity as a function of learning phases (one-way-ANOVA, F_2,38_ = 3.02, P = 0.06; Figure 5C-D).

**Figure 5:**
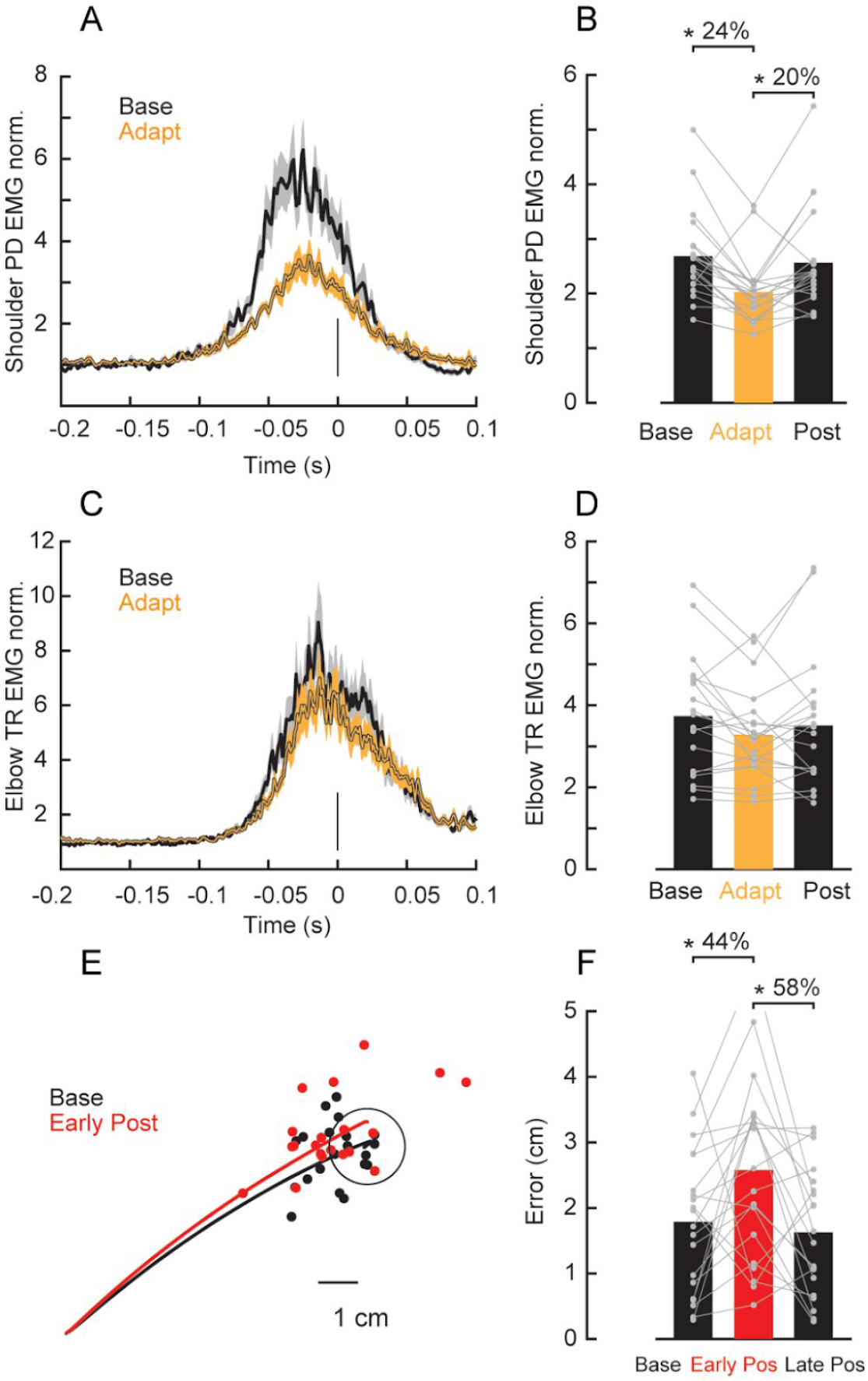
Transfer to reaching. **A**, Time series of shoulder extensor (PD) normalized muscle activity during elbow extension reaching trials averaged over the last 10 baseline and adaptation trials. Shaded areas represent the standard error of the mean. Data is aligned on movement onset. **B**, Average PD muscle activity in a fixed time window (−100 to 100 ms relative to movement onset) associated with trials late in the baseline, adaptation and post-adaptation. Each dot represents data from a single participant. Asterisks indicate reliable effects (p < 0.05, see main text). **C-D**, data for elbow TR muscle activity are shown using the same format as **A-B**. **E**, Average hand trajectories late in the baseline (10 trials) and early in the post-adaptation trials (first 3 trials). Each dot represents data from a single participant. F, Average error between hand position at movement offset to the center of the target in the last 10 trials in the baseline, first 3 trials early in the post-adaptation and last 10 trials late in post-adaptation phases (p < 0.05, see main text).

Additional evidence that learning of novel limb dynamics during perturbation trials transferred to reaching trials was the presence of kinematic after-effects. That is, in the early post-adaptation trials, participants generated trajectory errors in the direction predicted if they failed to compensate for the unlocked intersegmental dynamics (Figure 5E). We quantified these after-effects by performing a one-way ANOVA to compare reach accuracy (measured as distance from the center of the goal target) of trials late in the baseline phase (last 10 trials), trials early in the post-adaptation phase (first 3 trials) and trials late (last 10 trials) in the post-adaptation phase (Figure 5F). We chose a smaller bin size early in the post adaptation because the return to baseline after unlocking the shoulder joint happens very quickly (Maeda et al., 2018). We found a significant effect of phase on these trajectory errors (F_2,38_ = 5.53, P = 0.007). Tukey post-hoc tests showed that movement errors increased by 47% (p = 0.02) from baseline to early post-adaptation and returned to baseline levels (p = 0.86) in late post-adaptation trials.

### Learning and transfer with 100 ms load duration

In the experiments above we found that learning of arm dynamics during perturbation trials unfolded slowly over hundreds of trials and was ultimately incomplete, as shoulder muscle responses in the long-latency epoch only decreased by 47% from baseline trials rather than being completely eliminated. We also found an incomplete transfer to reaching trials as shoulder extensor activity decreased by only 24% from baseline reaching trials. To determine whether such limited learning and transfer arose because our paradigm yielded learning only in the long-latency reflex epoch, thirty additional participants performed the same experimental procedures as above except that the duration of the mechanical perturbation lasted 100 ms which evoked robust muscle activity in the so-called voluntary response epoch (>100 ms post-perturbation). After shoulder fixation, we observed changes in shoulder extensor activity both in the long-latency epoch (one-way-ANOVA, F_2,38_ = 14.09, P < 0.0001, and Tukey post-hoc tests showed that shoulder extensor muscle activity in the long-latency epoch decreased by 50% relative to baseline, P < 0.0001; Figure 6 A-B) and in the voluntary response (F_2,38_ = 7.19, P = 0.002, and Tukey post-hoc tests revealed that shoulder activity in this voluntary epoch decreased by 75% relative to baseline, P = 0.003; Figure 6 C-D). However, as for the 20 ms perturbations, such changes unfolded slowly, were incomplete and led to incomplete transfer to reaching, as shoulder extensor muscles during reaching decreased by 18.7% relative to baseline (one-way-ANOVA, F_2,38_ = 12.83, P < 0.0001, Tukey post-hoc tests showed that shoulder extensor muscle activity decreased relative to baseline trials, P < 0.0001; Figure 6 E-F).

**Figure 6:**
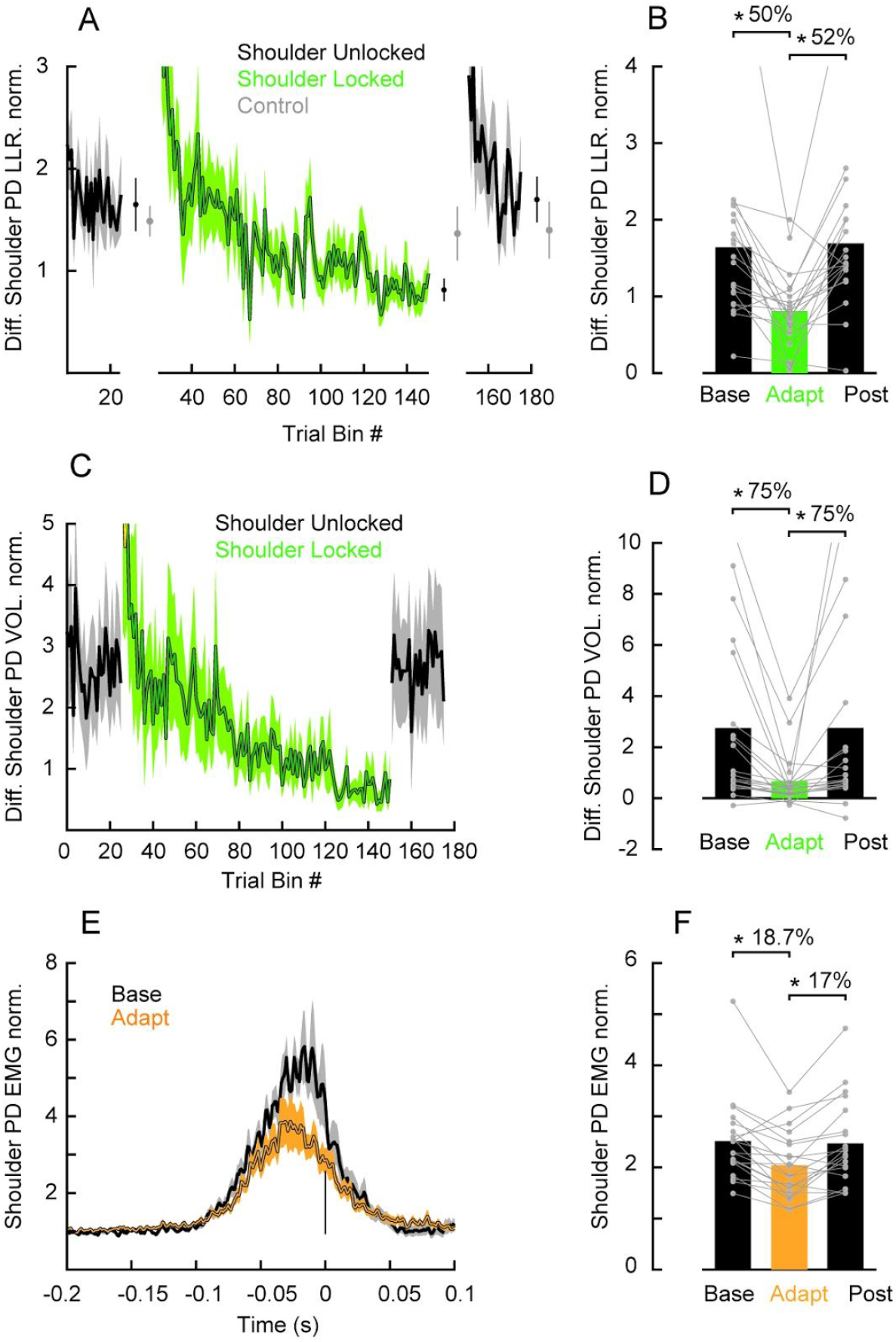
Learning novel arm dynamics during 100ms perturbation trials with shoulder fixation. **A**, Average of the difference of shoulder PD muscle activity (excitatory - inhibitory) in the long latency epoch (50-100ms post perturbation) across trials. Each data bin is 2 trials. Shaded areas represent the standard error of the mean. EMG data are normalized as described in the Methods. Error bars plotted between the phases represent the mean and standard error of the last 5 bins of trials in each phase contrasted with the respective bins in the control experiment in gray. **B**, Average PD muscle activity in the long-latency epoch associated with trials late in the baseline, adaptation, and post-adaptation. Each dot represents data from a single participant. **C-D**, data for the difference in the voluntary epoch (excitatory-inhibitory) are shown using the same format as **A-B**. **E**, Time series of shoulder PD normalized muscle activity during elbow extension reaching trials averaged over the last 10 baseline and adaptation trials. Shaded areas represent the standard error of the mean. Data is aligned on movement onset. **F**, Average PD muscle activity in a fixed time window (−100 to 100 ms relative to movement onset) associated with trials late in the baseline, adaptation and post-adaptation. Each dot represents data from a single participant. Asterisks indicate reliable effects (P < 0.05, see main text).

## Discussion

Here we tested whether the nervous system learns novel intersegmental dynamics following shoulder fixation during posture perturbations and probed whether such learning transfers to self-initiated reaching. We did so by applying 20 ms perturbations that created pure elbow motion with the shoulder joint free and fixed, while asking participants to not intervene with the perturbation. We found that shoulder muscle responses in the long-latency epoch (50-100ms post perturbation) slowly decreased after the shoulder was mechanically fixed by the robotic manipulandum. After this learning phase, we found that agonist shoulder muscle activity during self-initiated elbow extension reaching trials also decreased and participants showed increased kinematic errors in the direction expected if they failed to properly compensate for the mechanical effect of forearm rotation at the shoulder joint. Taken together, our results demonstrate that the nervous system learns novel arm dynamics during feedback control and that this learning transfers from feedback to feedforward control.

### Internal models for feedback control

Internal models are a well-established concept in self-initiated reaching (Wolpert et al., 1995). Internal models allow voluntary control mechanisms to compensate for the complex mechanical properties of the arm, delayed and noisy sensory feedback and to adapt to changes of the body or the environment (Kawato, 1999; Wolpert and Flanagan, 2001). Less appreciated is that internal models also apply to fast feedback responses, which have to compensate for the same factors when countering external perturbations (Lacquaniti and Soechting, 1986; Gielen et al., 1988; Kurtzer et al., 2008, 2009, 2014; Wagner and Smith, 2008; Pruszynski et al., 2011; Ahmadi-Pajouh et al., 2012; Cluff and Scott, 2013; Crevecoeur and Scott, 2013, 2014; Weiler et al., 2016; Maeda et al., 2017, 2018; Kurtzer, 2019). For example, an internal model of the arm enables long-latency stretch reflexes, a class of fast feedback responses sensitive to mechanical stretch that include a cortical neural contribution (Pruszynski and Scott, 2012; Scott, 2016), to account for the arm’s intersegmental dynamics and respond appropriately to the applied joint torques rather than local joint motion (Lacquaniti and Soechting, 1986; Kurtzer et al., 2008, 2009, 2014; Maeda et al., 2017, 2018; Kurtzer, 2019).

There are two main findings in this study. First, that long-latency stretch reflexes can directly update the internal model that maps joint motion to joint torque without explicitly engaging voluntary control mechanisms. Second, that the updated internal model learned by long-latency stretch reflexes applies during self-initiated reaching. These findings add to a growing body of work demonstrating how internal models updated during self-initiated reaching transfer to fast feedback control (Wagner and Smith, 2008; Ahmadi-Pajouh et al., 2012; Yousif and Diedrichsen, 2012; Cluff and Scott, 2013; Maeda et al., 2018) by showing that such learning and transfer is bidirectional. These findings are also consistent with modern theories of motor control, based on optimal feedback control, which posit that motor behavior is achieved via the sophisticated manipulation of sensory feedback (Todorov and Jordan, 2002; Scott, 2004). Under this class of models, bidirectional transfer between feedforward and feedback control is expected because feedforward motor commands and transcortical feedback responses are part of the same control system implemented in common neural circuits (Pruszynski, 2014; Scott, 2016).

An important avenue of future research is determining which neural circuits underlie shared learning during feedforward and feedback control (for review, see Scott, 2016). One likely common node is the primary motor cortex. Previous work has shown that neurons in primary motor cortex are engaged during both self-initiated reaching actions and following mechanical perturbations applied to the arm (Evarts, 1973; Evarts and Tanji, 1976; Wolpaw, 1980; Evarts and Fromm, 1981; Picard and Smith, 1992; Li et al., 2001; Gribble and Scott, 2002; Pruszynski et al., 2011, 2014; Omrani et al., 2014; Heming et al., 2016). In addition, recent studies have demonstrated that primary motor cortex is causally involved in the compensation for the arm’s intersegmental dynamics during both self-initiated reaching actions (Gritsenko et al., 2011) and in the context of externally applied mechanical perturbations (Pruszynski et al., 2011). Primary motor cortex activity is also modified during motor learning in the context of self-initiated reaching (Sanes and Donoghue, 2000; Li et al., 2001; Diedrichsen et al., 2005; Kawai et al., 2015), though, to our knowledge, no studies have definitively linked primary motor cortex to the learning process itself. Another likely common node, especially for the learning process, is the cerebellum, which is richly interconnected with primary motor cortex (Wagner et al., 2019). Cerebellar circuits are strongly implicated in multi-joint coordination during both feedforward control and feedback responses (Holmes, 1939; Goodkin et al., 1993; Bastian et al., 1996, 2000; Kurtzer et al., 2013; Wagner et al., 2019) and have long been hypothesized to house the computations related to internal models (Wolpert et al., 1998; Kawato, 1999).

It is also possible that the learning itself, while mediated via the cortical or cerebellar structures, is implemented at the level of the spinal cord as both feedforward and feedback motor commands must ultimately pass through spinal interneurons and motorneurons projecting to the muscle. In an elegant set of experiments, Wolpaw and colleagues have studied the operant conditioning of the H-reflex, the electrically-induced analog of the spinal stretch reflex, and its contribution to rehabilitation following spinal cord injury (for review, see Thompson and Wolpaw, 2014). They have shown that such conditioning produces multisite changes at the level of the spinal cord that actually drive the observed differences in the H-reflex response, including a shift in motorneuron firing threshold (Carp and Wolpaw, 1994) and a change in the number of GABAergic terminals (Wang et al., 2006). Importantly, successful operant conditioning of the spinal cord circuit itself requires a functional corticospinal tract and sensorimotor cortex as well as the cerebellum and inferior olive but no other major ascending or descending spinal pathways (Chen and Wolpaw, 2002, 2005; Chen et al., 2002, 2006a, 2006b, 2016; Wolpaw and Chen, 2006), indicating that cerebellar contributions via the sensorimotor cortex (as opposed to the rubrospinal tract) are critical for implementing the learning (Chen and Wolpaw, 2005; Wolpaw and Chen, 2006). Given the differences between H-reflex operant conditioning, especially with respect to its development over weeks and months, an extremely long time-scale even relative to the slow learning in our paradigm, it is unclear whether the same mechanisms are in play for the type or learning we report here. However, at the very least, the general concept and experimental approach serves a useful roadmap for examining the plastic changes associated with learning new intersegmental dynamics and how such learning is commonly implemented by feedforward and feedback control systems.

### Limitations

One goal of our experimental design was to minimize or eliminate the engagement of voluntary responses to mechanical perturbation in the learning process so that we could attribute any observed learning directly to the neural mechanisms that generate feedback responses rather than learning from whatever mechanisms generate voluntary motor commands. We did this by using very short duration perturbations (20 ms) and instructing participants to not intervene with the perturbations, both of which have been previously shown elicit long-latency stretch reflexes (50-100 ms post-perturbation) while reducing or eliminating associated voluntary responses (>100 ms post-perturbation) (Asatryan and Feldman, 1965; Crago et al., 1976; Ghez and Shinoda, 1978; Lee and Tatton, 1982; Calancie and Bawa, 1985; Lewis et al., 2005; Pruszynski et al., 2008; Schuurmans et al., 2009; Shemmell et al., 2009; Kurtzer et al., 2010; Forgaard et al., 2015, 2016, 2019; Kurtzer, 2019). Our paradigm did create the expected response profiles such as no reliable decrease in muscle activity after 100 ms post-perturbation with shoulder fixation, presumably because the evoked response in the voluntary epoch was minimal to begin with (Figure 4 A-B). However, it is important to emphasize that we cannot establish that participants did not engage self-initiated responses as we have no direct or independent measurement of these responses.

## Acknowledgements

We thank Chris Forgaard for comments on the manuscript.

